# Enabling complex fibre geometries using 3D printed axon-mimetic phantoms

**DOI:** 10.1101/2021.12.07.471599

**Authors:** Tristan K. Kuehn, Farah N. Mushtaha, Ali R. Khan, Corey A. Baron

**Author notes:** These authors share first authorship (Ali R. Khan, Corey A. Baron). These authors share senior authorship. **Contributions to the field** Diffusion magnetic resonance imaging is sensitive to the random motion of water molecules. Axonal tracts in the brain restrict this motion, making the signal measured by diffusion magnetic resonance imaging sensitive to the orientation of the axons. To better interpret this signal in the brain, mathematical models or representations can be applied to the measured signal. The underlying fibre structure in the brain is unknown, so MRI phantoms with known fibre structure can be used to characterize a model or representation’s performance. Here we introduce a new method for producing 3D printed axon-mimetic phantoms with bending, fanning, and kissing fibre structures, which are challenging for models and representations to accurately describe. We use a set of these phantoms to characterize the performance of several models and signal representations in the presence of these fibre structures. The phantoms allow physically-acquired scan data to be used for characterizing model and representation performance easily and inexpensively. We use the phantoms to identify a set of metrics that appear to robustly indicate microstructural change in the presence of orientation dispersion of anisotropic fibres, and confirm that a set of metrics designed to indicate orientation dispersion do so.

## Abstract

**Purpose:** To introduce a method to create 3D-printed axon-mimetic phantoms with complex fibre orientations to characterize the performance of diffusion MRI models and representations in the presence of orientation dispersion.

**Methods:** An extension to an open source 3D printing package was created to 3D print a set of five 3D-printed axon-mimetic (3AM) phantoms with various combinations of bending and crossing fibre orientations. A two-shell diffusion MRI scan of the five phantoms in water was performed at 9.4T. Diffusion tensor imaging (DTI), diffusion kurtosis imaging (DKI), the ball and stick model, neurite orientation density and dispersion imaging (NODDI), and Bingham-NODDI were all fit to the resulting diffusion MRI data. A fiducial in each phantom was used to register a ground truth map of that phantom’s crossing angles and/or arc radius to the diffusion-weighted images. Metrics from each model and representation were compared to the ground-truth maps, and a quadratic regression model was fit to each combination of output metric and ground-truth metric.

**Results:** The mean diffusivity (MD) metric defined by DTI was insensitive to crossing angle, but increased with fibre curvature. Axial diffusivity (AD) decreased sharply with increasing crossing angle. DKI’s diffusivity metrics replicated the trends seen in DTI, and its mean kurtosis (MK) metric, decreased with fibre curvature, except in regions with high crossing angles. The estimated stick volume fraction in the ball and stick model decreased with increasing fibre curvature and crossing angle. NODDI’s intra-neurite volume fraction was insensitive to crossing angle, and its orientation dispersion index (ODI) was strongly correlated to crossing angle. Bingham-NODDI’s intra-neurite volume fraction was also insensitive to crossing angle, while its primary ODI (ODI_P_) was also strongly correlated to crossing angle and its secondary ODI (ODI_S_) was insensitive to crossing angle. For both NODDI models, the volume fractions of the extra-neurite and CSF compartments had low reliability with no clear relationship to crossing angle.

**Conclusions:** This study demonstrates that inexpensive 3D-printed axon-mimetic phantoms can be used to investigate the effect of fibre curvature and crossings on diffusion MRI representations and models of diffusion signal. As a proof of concept, the dependence of several representations and models on fibre dispersion/crossing were investigated. As expected, Bingham-NODDI was best able to characterize planar fibre dispersion in the phantoms.

## 1 Introduction

Diffusion MRI (dMRI) is an imaging modality that is sensitive to the diffusion of water molecules on a microscopic scale. The dMRI signal in a voxel containing one or more axonal fibres is determined by both the microstructure and the orientation of the fibres. Diffusion MRI representations and models of white matter characterize microstructure and orientation with varying levels of sensitivity and specificity. Discriminating between signal variation caused by microstructural changes and signal variation caused by orientational changes is difficult, and dMRI models are known to characterize microstructure less precisely in the presence of orientation dispersion, where fibre segments with different orientations within a single voxel cause partial volume errors (Alexander et al., 2001). These complex fibre configurations are widespread in the human brain (Jeurissen et al., 2013), so the ideal dMRI model of white matter would characterize both microstructure and orientation accurately and robustly.

To validate dMRI models of white matter, it would be useful to quantify the effect of orientation dispersion on those models’ estimated parameters. However, it is difficult to perform that quantification because there is no widely accepted complementary modality to provide a ground truth of axonal orientation *in vivo*. Instead, numerical or physical phantoms can provide a ground truth (Fieremans and Lee, 2018). Numerical phantoms allow precise control of the sample, but simulating a dMRI scan of an anatomically realistic sample is computationally intensive, limiting the feasible volume of a region of interest. Physical phantoms offer real scan data with a ground truth. However, existing physical fibre-containing phantoms tend to be expensive and/or time-consuming to prepare, with limited orientational complexity.

Fused deposition modeling (FDM) 3D printing offers a promising technology for producing dMRI phantoms, if an appropriate material is used. GEL-LAY (LAY Filaments, Cologne, Germany) is an FDM filament composed of an elastomeric matrix containing pockets of polyvinyl alcohol (PVA). When 3D printed, the pockets of PVA form fibres with a long axis oriented along the direction of motion of the print head. When immersed in water, the PVA dissolves, leaving microscopic fibrous pores that can mimic the diffusion characteristics of axonal fibres, with known primary directions of diffusion. The diameters of these pores follow a gamma distribution with a median diameter of 3.5 μm (Mushtaha et al., 2021). These 3D printed axon-mimetic (3AM) phantoms are inexpensive and effective, but have not yet been used to produce a phantom with complex configurations of crossing fibres.

Here, we introduce a method to create bending, kissing, and fanning patterns of crossing fibres. We then use phantoms 3D printed with this method to characterize the effect of those complex geometries on several well-known dMRI models.

## 2 Materials and Methods

### 2.1 Open-source 3D printing extension

A custom extension to the Ultimaker Cura 3D printer software was developed to allow within-layer fibre configurations composed of concentric arcs to be produced (https://github.com/tkkuehn/CuraEngine), in addition to the patterns composed of parallel lines available by default.

### 2.2 3D printed phantoms

We used the Cura extension to design and 3D print five phantoms (Figure 1), and prepared the phantoms according to a protocol available in the Supporting information and hosted at <u>osf.io/zrsp6</u> (Mushtaha et al., 2021). The phantoms were cylindrical, with a radius of 11 mm and height of 4.5 mm. Each phantom was composed of 45 0.1 mm thick layers of printed material.

**Figure 1.**
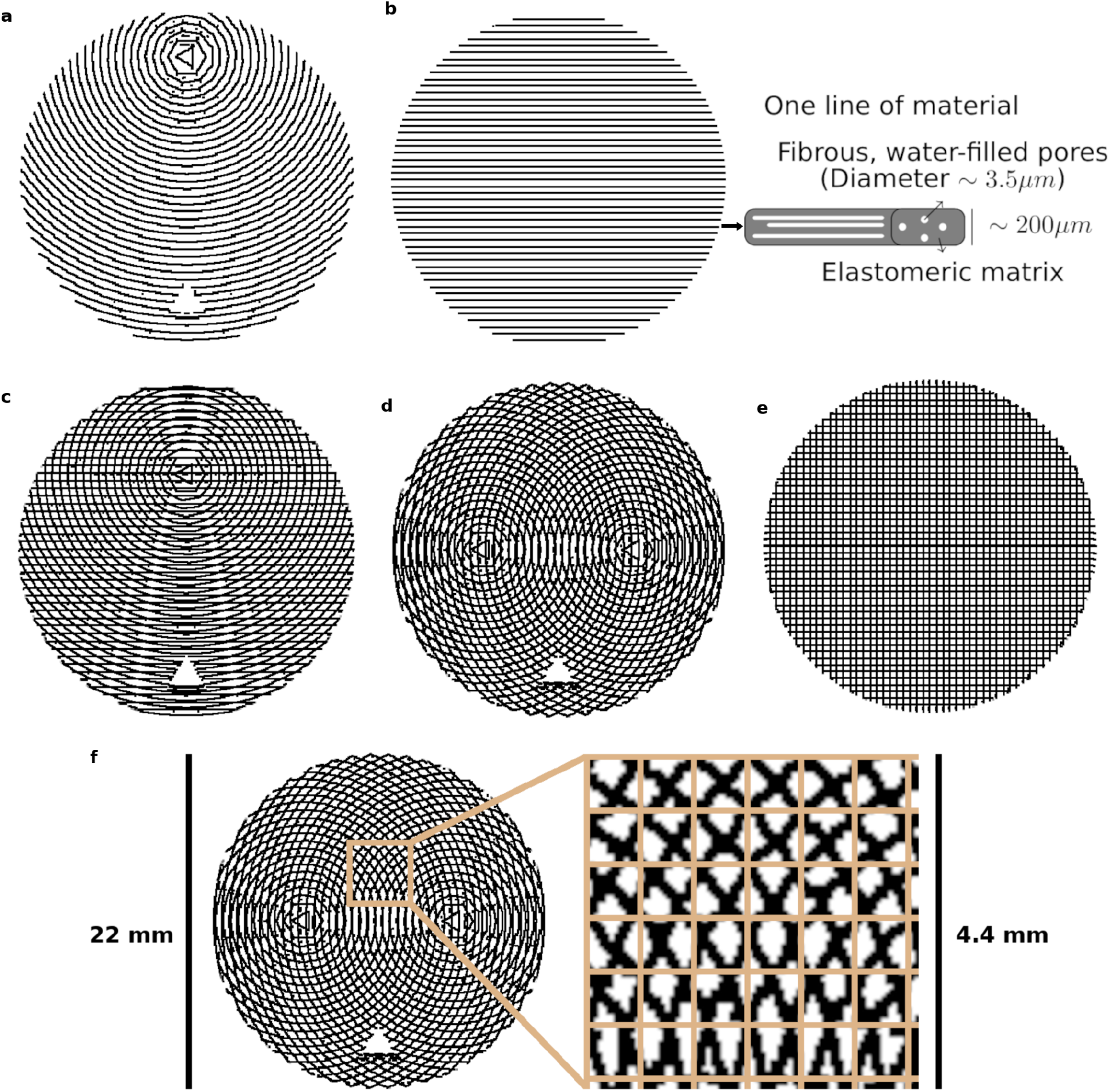
The five phantom patterns, with alternating layers superimposed on one another. (**A**) Bending. (**B**) Straight, with a schematic of one line of material. (**C**) Kissing. (**D**) Fanning. (**E**) Crossing (**F**) Crossing, with the isotropic in-plane voxel size overlaid on a cut-out.

Each phantom was designed with a different fibre configuration, achieved by changing the pattern in which lines of material were deposed in each layer. Phantoms intended to mimic interacting axonal tracts were created by alternating between two different patterns from layer to layer. The five phantoms were produced to mimic bending, straight, kissing, fanning, and crossing fibres, as described in Table 1 and illustrated in Figure 1. The three phantoms with spatially heterogeneous fibre configurations (i.e. the bending, fanning, and kissing phantoms) were printed with a triangular hole to serve as a fiducial.

**Table 1.** Description of the within-layer fibre configurations of each phantom. Phantoms with two layer patterns alternate between those patterns from layer to layer.

| Label | Layer pattern 1 | Layer pattern 2 |
| --- | --- | --- |
| Bending | Concentric arcs with a centre 10 mm from the phantom's axis. | N/A |
| Straight | Parallel lines. | N/A |
| Kissing | Parallel lines. | Concentric arcs with a centre 6.5 mm from the phantom's axis. |
| Fanning | Concentric arcs with a centre 5 mm from the phantom's axis. | Concentric arcs with a centre 10 mm away from the other pattern's centre, directly opposite. |
| Crossing | Parallel lines. | Parallel lines, perpendicular to the lines in |

**Table 2.** List of metrics from each model assessed in relationship to ground truth phantom metrics.

| Model or Representation | Metrics | References |
| --- | --- | --- |
| DTI | AD, RD, MD, FA | (Basser et al., 1994) |
| DKI | AD, RD, MD, FA, AK, RK, MK | (Jensen et al., 2005; Lu et al., 2006) |
| Ball and Stick | Stick volume fraction, Ball volume fraction | (Behrens and Woolrich, 2003) |
| NODDI | Intra-neurite volume fraction, Extra-neurite volume fraction, CSF volume fraction, ODI | (Zhang et al., 2012) |
| Bingham-NODDI | Intra-neurite volume fraction, Extra-neurite volume fraction, CSF volume fraction, $ODI_P$, $ODI_S$ | (Tariq et al., 2016) |

### 2.3 MRI

Diffusion MRI was implemented with a 9.4 T Bruker small animal scanner using using a single-shot EPI sequence with 120 and 60 directions at b=2000 and 1000 s/mm^2^, respectively, 20 averages at b=0 s/mm^2^, diffusion gradient lobe duration (δ) of 4.1 ms, spacing between gradient lobes (Δ) of 13.1 ms, gradient magnitudes calculated to achieve the intended b-values, TE/TR=37/2500 ms, FOV=200×200 mm^2^, 0.7 mm isotropic in-plane resolution, and one 3 mm axial slice per phantom.

### 2.4 Analysis

For the fanning, bending, and crossing phantoms, we localized the phantom’s centroid and fiducial in the mean Diffusion Weighted Image (DWI), and used those points to produce ground truth images of crossing angle or radius of curvature, as appropriate, registered to the MRI images (Figure 2).

**Figure 2:**
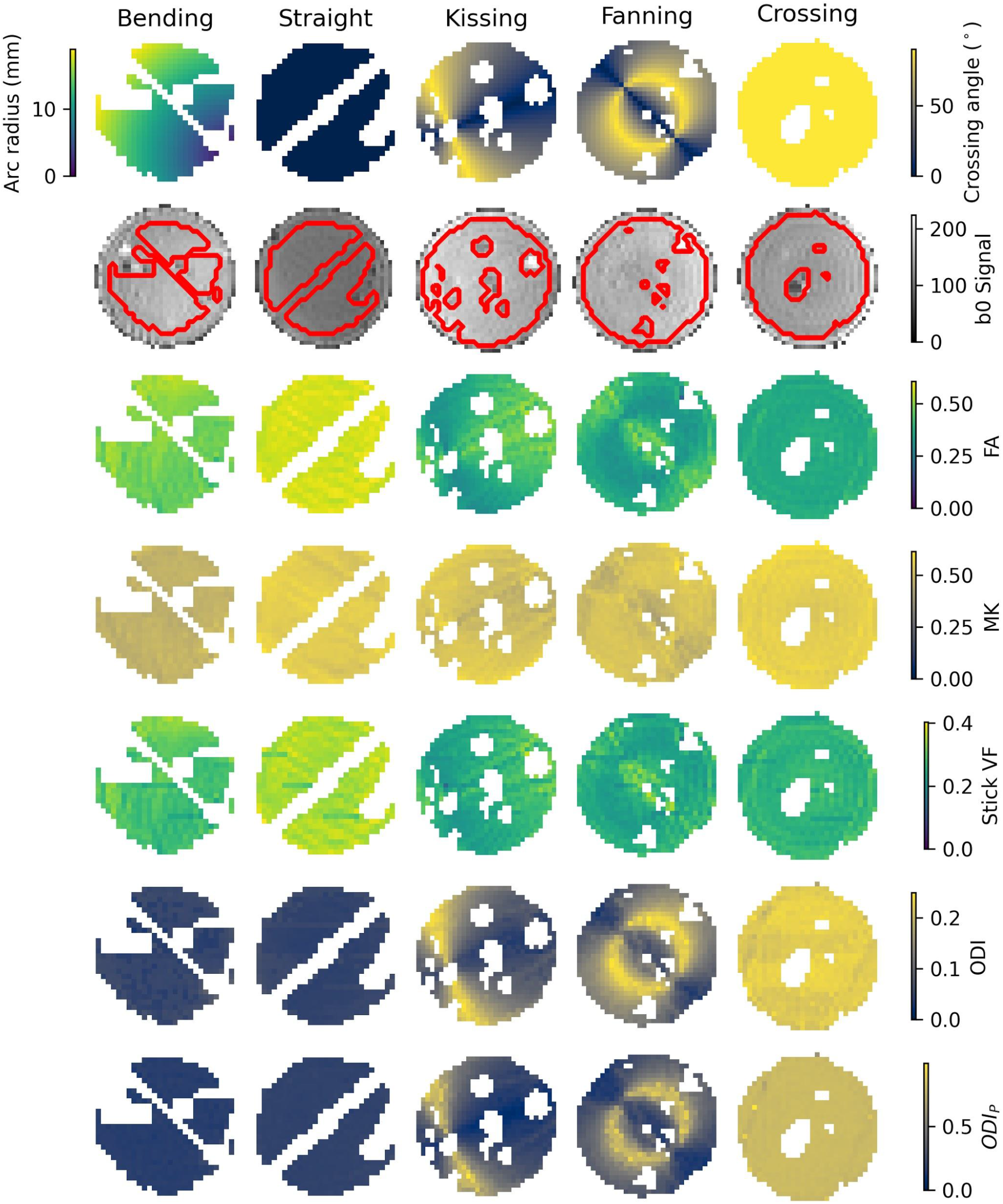
Parameter maps of each phantom. Top row: Ground truth geometry metrics (Arc radius in the bending phantom, crossing angle in the others). Second row: b0 images with a contour of the mask superimposed. Third row: FA. Fourth row: MK. Fifth row: Stick volume fraction. Sixth row: ODI. Seventh row: ODI_P_.

DIPY (Garyfallidis et al., 2014) was used to perform weighted ordinary least squares fits of Diffusion Tensor Imaging (DTI) (Basser et al., 1994), using only the acquisitions at b = 0 and 1000 s/mm^2^; and Diffusion Kurtosis Imaging (DKI) (Jensen et al., 2005; Lu et al., 2006) representations to the DWI’s. Four parameters were then extracted from the DTI fit: axial diffusivity (AD), which quantifies the estimated diffusivity along a central axis; radial diffusivity (RD), which quantifies the diffusivity in the directions perpendicular to that central axis; mean diffusivity (MD), which quantifies the mean diffusivity across all directions; and fractional anisotropy (FA), which quantifies the degree to which diffusion in a voxel is anisotropic. The same four diffusivity-based metrics are extracted from the DKI, in addition to three kurtosis metrics, which capture the degree to which diffusion in a voxel deviates from Gaussianity: axial kurtosis (AK), radial kurtosis (RK), and mean kurtosis (MK).

The Microstructure Diffusion Toolbox (MDT) (Harms et al., 2017) was used to perform Powell conjugate-direction optimized fits of ball and stick (Behrens and Woolrich, 2003), NODDI (Zhang et al., 2012), and Bingham-NODDI (Tariq et al., 2016) models to the DWI. All models’ assumed diffusivities were fixed at 2.2 mm^2^/s, the diffusivity of pure water at room temperature, based on the DTI diffusivity metrics observed in a straight phantom (Mushtaha et al., 2021). The single parameter extracted from the ball and stick model is the relative volume fraction between the ball compartment and the stick compartment. NODDI and Bingham-NODDI both produce volume fractions of intra-neurite, extra-neurite, and CSF compartments. NODDI also produces the orientation dispersion index (ODI), which quantifies from 0-1 the dispersion of the Watson distribution that governs the intra- and extra-neurite compartment. Bingham-NODDI splits ODI into a primary ODI (ODI_P_), and a secondary ODI (ODI_S_), quantifying the anisotropic dispersion described by the Bingham distribution.

For each dMRI representation or model parameter, we fit three quadratic regression models: the parameter vs. radius of curvature in the bending phantom, and the parameter vs. crossing angle in the kissing and fanning phantoms. The R^2^ was used to quantify how much dMRI model variance was accounted for by each regression model.

## 3 Results

An example b=0 s/mm^2^ image, registered ground truth map, and metric map from each model or representation is shown for each phantom in Figure 2. To omit regions with air bubbles, a mask for each phantom was manually drawn using the b0 images as a guide. Overall, strong correspondence is observed between the ground truth fibre geometries and the quantitative dMRI maps. Quantitative comparisons for each of the representations and models will be described below.

### 3.1 DTI

Figure 3 plots the DTI metrics in each phantom against metrics related to the orientation dispersion in that phantom. The DTI metrics in the straight phantom describe a very anisotropic diffusion signal with AD ∼ 2.2 mm^2^/s and RD ∼ 1.0 mm^2^/s, leading to an FA of about 0.55. MD was the only DTI metric that showed no relationship with crossing, but did show a relationship with arc radius in the bending phantom. Conversely, AD and FA were particularly strongly correlated to crossing angle.

**Figure 3:**
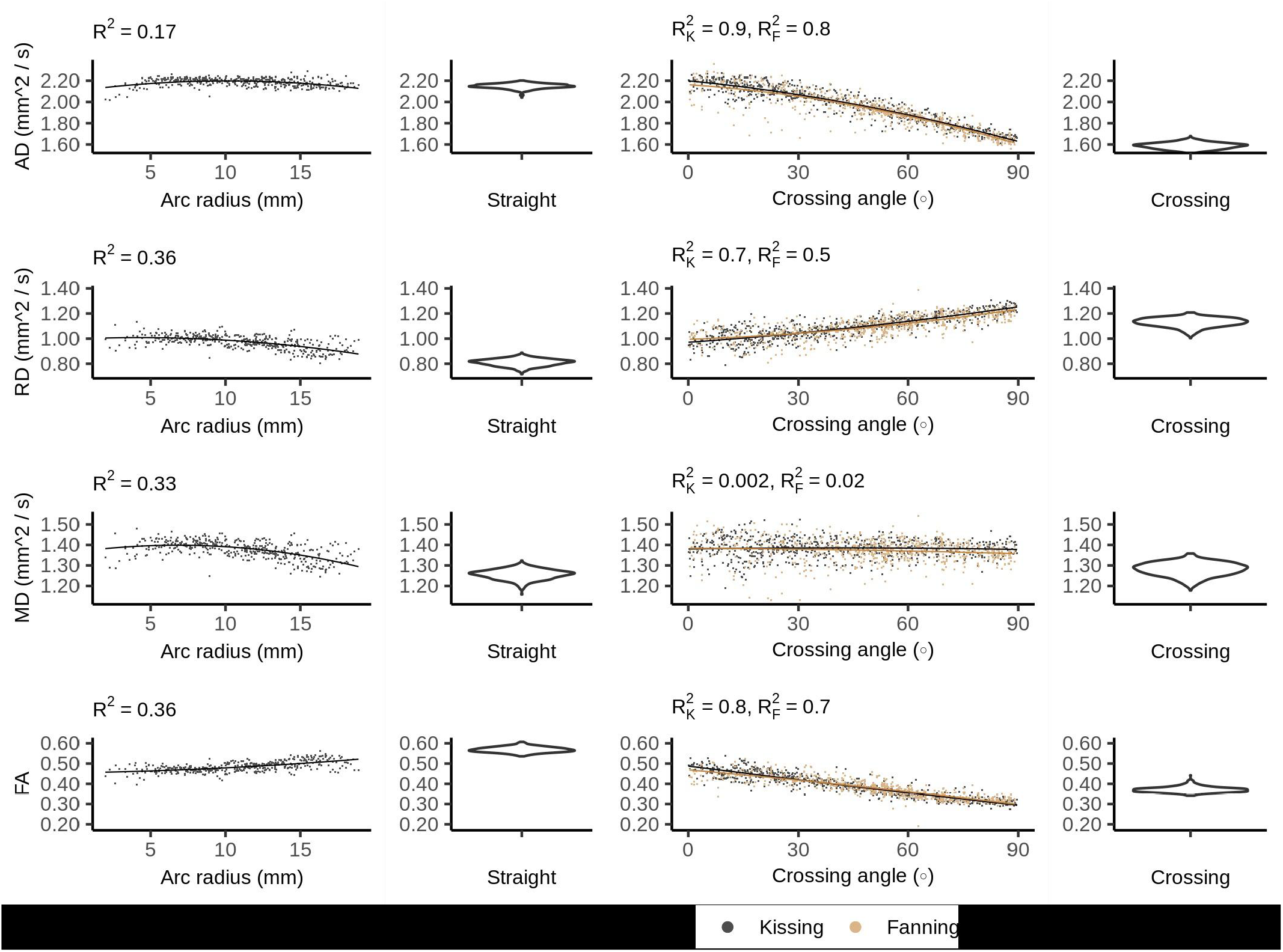
DTI metrics vs. orientation dispersion metrics in each of the four phantoms. Each sample corresponds to a single voxel. Each row depicts the relationship between one metric and orientation dispersion, and each column depicts the data from one or two phantoms. From left to right, those phantoms are: bending, straight, fanning and kissing, and crossing.

### 3.2 DKI

The DKI diffusivity metrics (AD, RD, MD, and FA) show similar patterns to those observed in DTI, but with less pronounced relationships to arc radius and crossing angle (Figure 4).

**Figure 4:**
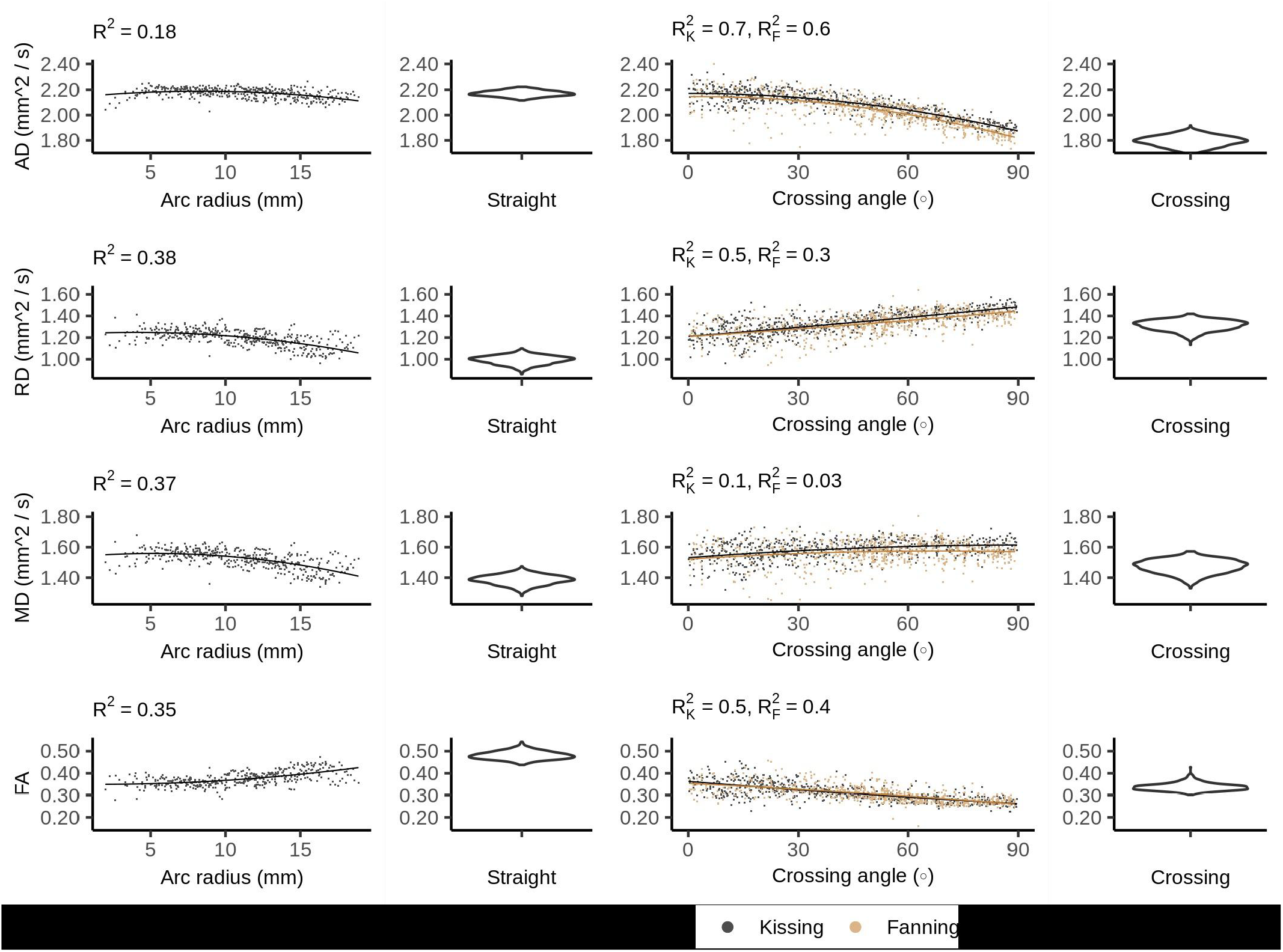
DKI diffusivity metrics vs. orientation dispersion metrics in each of the four phantoms. Each row depicts the relationship between one metric and orientation dispersion, and each column depicts the data from one or two phantoms. From left to right, those phantoms are: bending, straight, fanning and kissing, and crossing.

In the straight phantom there is little axial kurtosis (AK) and a median radial kurtosis (RK) of about 1.15 (Figure 5). Those metrics combine to produce a mean kurtosis ∼0.55. Arc radius explains little of the variance in the kurtosis metrics. The mean value and variance of AK both increase at higher fibre crossing angles, a pattern that is not observed in any other metric. There is little difference in the distribution of MK between the straight and crossing phantoms, but MK increases with crossing angle in the fanning and kissing phantoms.

**Figure 5:**
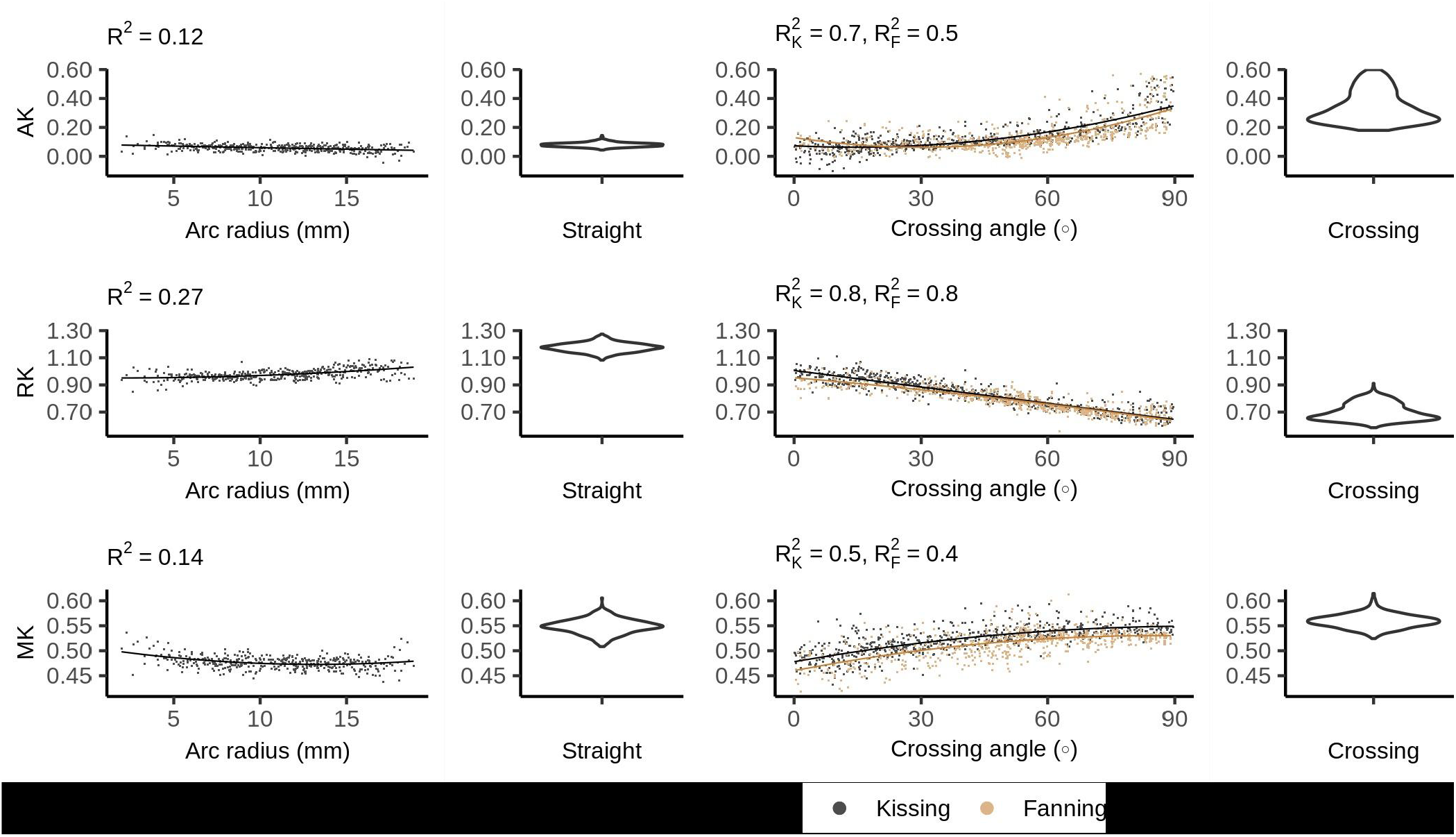
DKI kurtosis metrics vs. orientation dispersion metrics in each of the four phantoms. Each row depicts the relationship between one metric and orientation dispersion, and each column depicts the data from one or two phantoms. From left to right, those phantoms are: bending, straight, fanning and kissing, and crossing.

### 3.3 Ball and Stick

Figure 6 shows the ball and stick volume fractions in each phantom. The straight phantom is assigned a stick volume fraction of about 0.32, roughly half the ball volume fraction. The stick volume fraction increases with arc radius, and decreases with crossing angle.

**Figure 6:**
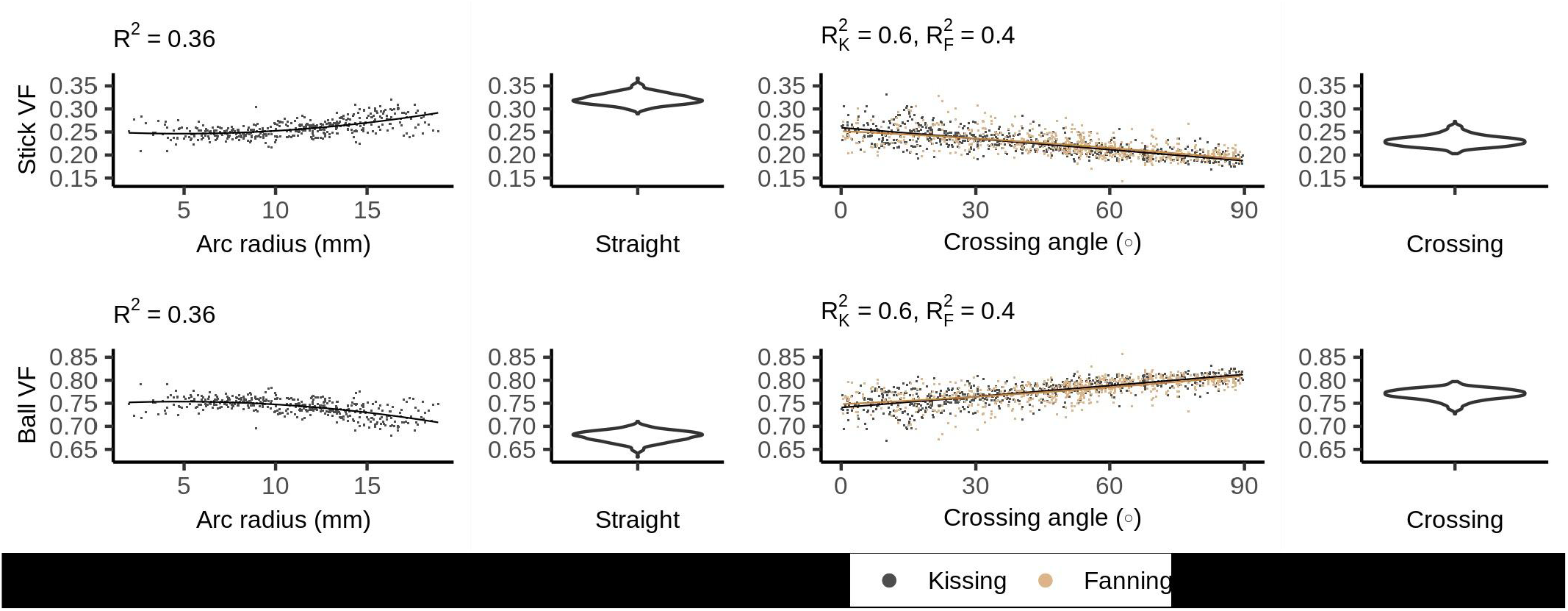
Ball and stick volume fractions vs. orientation dispersion metrics in each of the four phantoms.

### 3.4 NODDI

Figure 7 shows the NODDI parameters for each phantom. In all phantoms, there is a bimodal distribution of volume fraction assigned to the CSF and the extra-neurite compartments, while the intra-neurite volume fraction is stable. Intra-neurite volume fraction increases at higher arc radii, while the other volume fractions are not strongly related to arc radius or crossing angle. ODI is tightly related to crossing angle, spanning from 0 to about 0.25.

**Figure 7:**
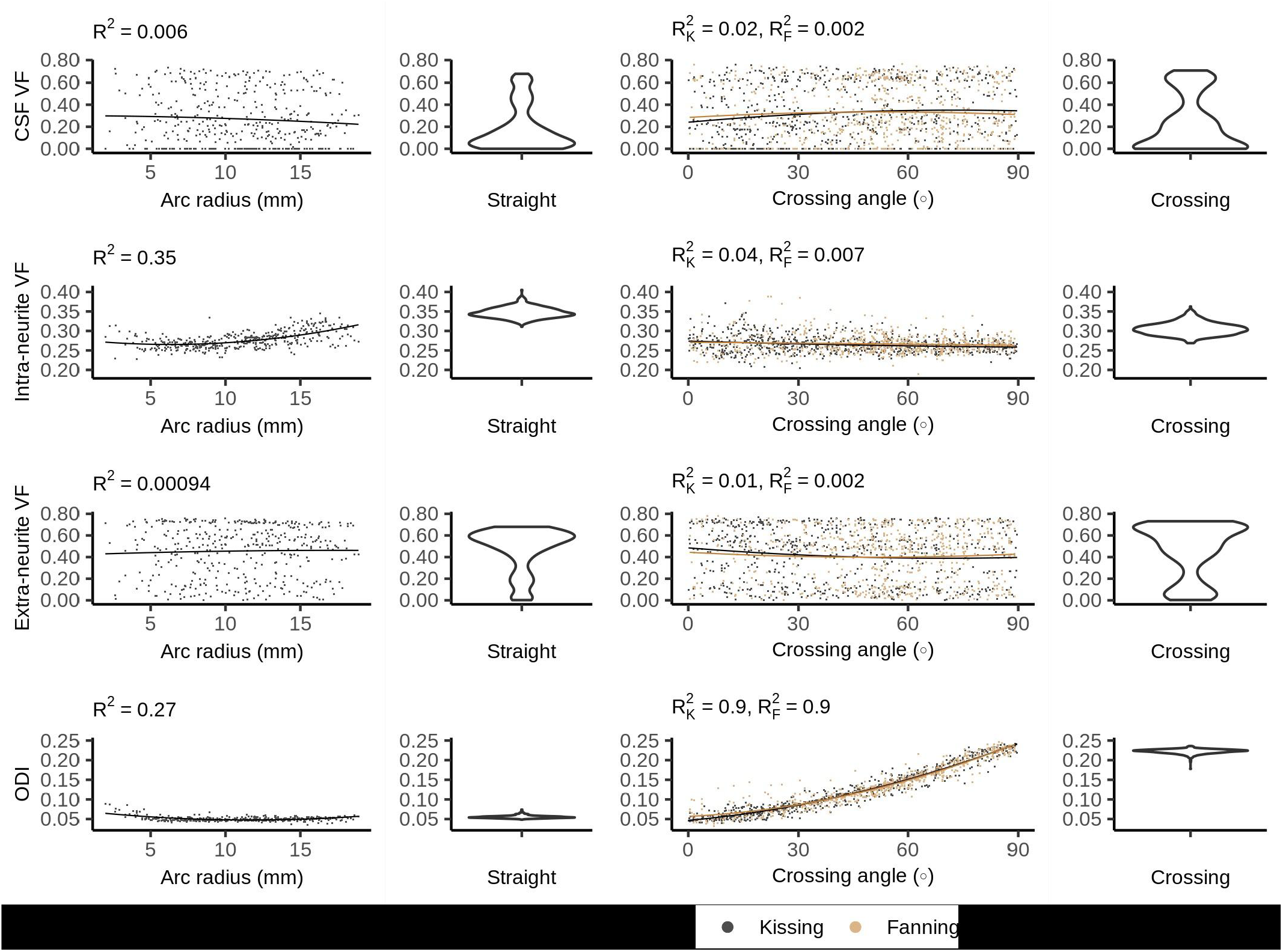
NODDI metrics vs. orientation dispersion metrics in each of the four phantoms.

### 3.5 Bingham-NODDI

Figure 8 plots the Bingham-NODDI parameters in each phantom against the orientation dispersion in that phantom. Figure 9 maps Bingham-NODDI’s dominant orientation, primary dispersion orientation, and secondary dispersion orientation in each phantom.

**Figure 8:**
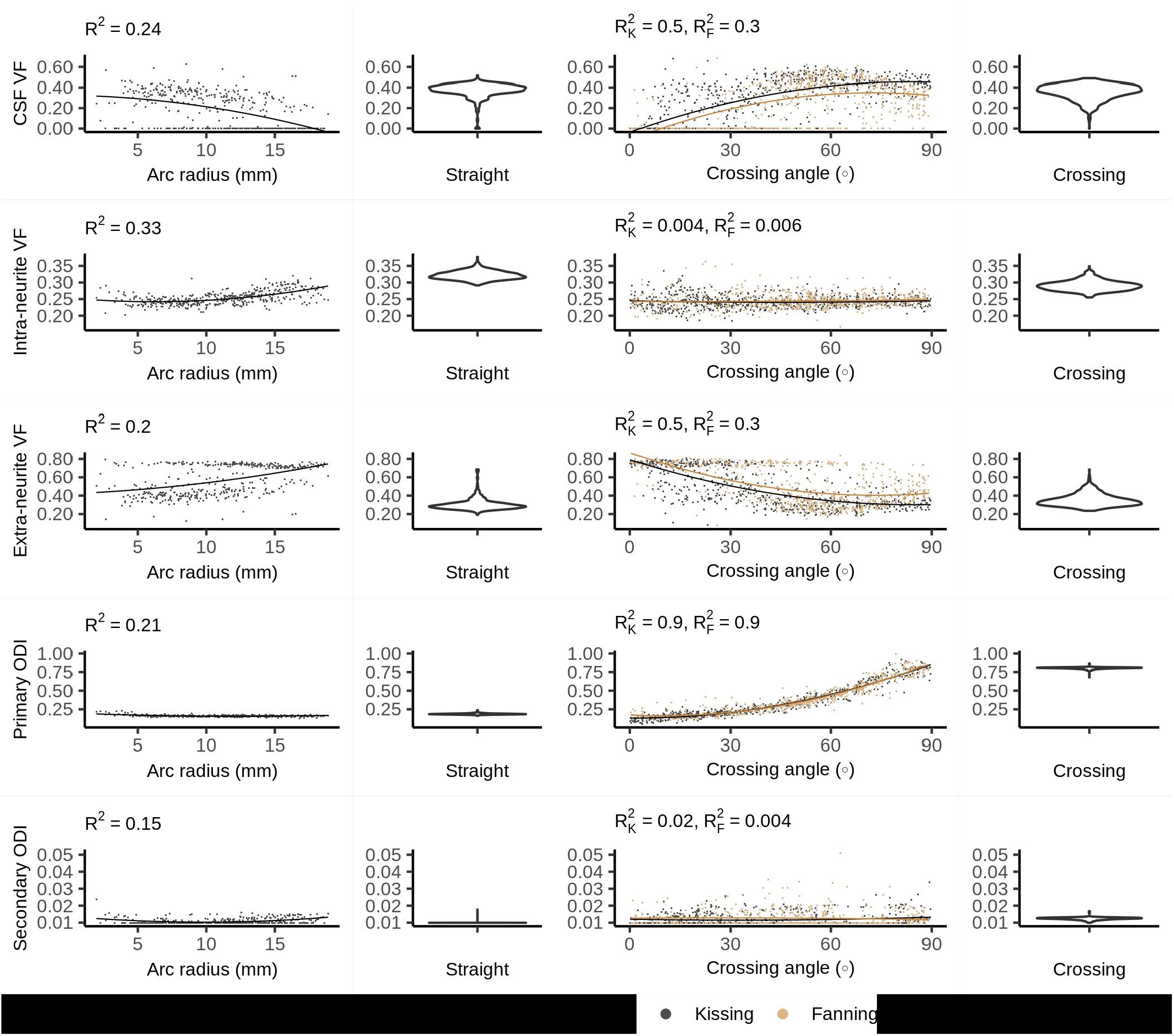
Bingham-NODDI metrics vs. orientation dispersion metrics in each of the four phantoms.

**Figure 9:**
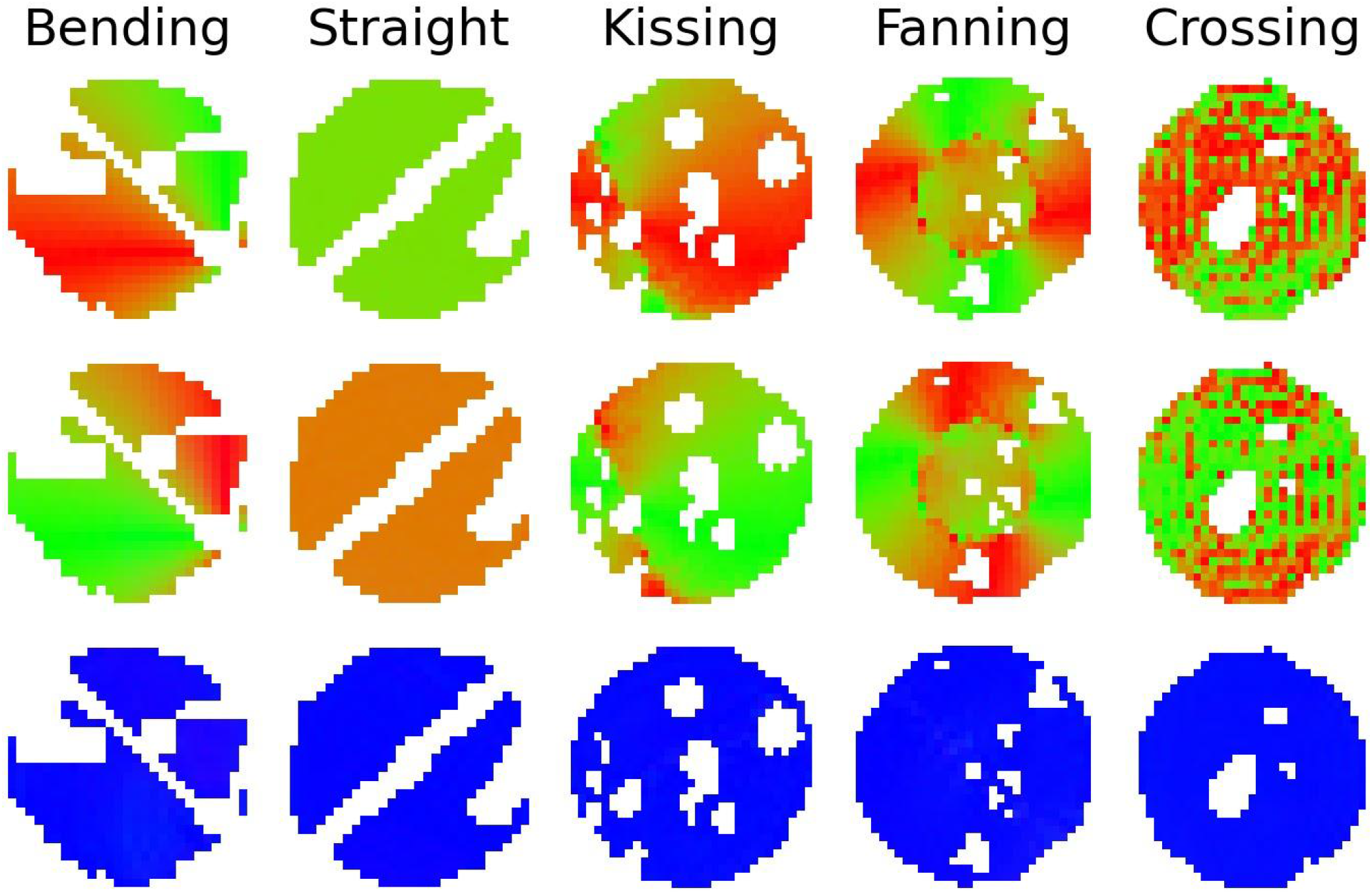
Colour maps of the three orthogonal orientations defined by Bingham-NODDI in each phantom. Top row: The dominant orientation. Middle row: The primary dispersion orientation. Bottom row: The secondary dispersion orientation.

Like NODDI, Bingham-NODDI’s intra-neurite volume fraction is stable, while the volume fraction assigned to the CSF and extra-neurite compartments is unstable: in most voxels, the CSF volume fraction is either zero or close to 0.4. Unlike NODDI, the split has a dominant mode with higher CSF volume fractions (and lower extra-neurite volume fractions), and appears to be biased by orientation dispersion: at high crossing angles and low radii of curvature, volume fraction appears more likely to be assigned to the CSF compartment. ODI_P_ is tightly related to crossing angle, and spans the defined range of the metric, between zero and one. ODI_S_, conversely, is not related to arc radius or crossing angle, and is below 0.05 in all voxels.

The dominant orientation and primary dispersion orientations are both restricted to the printing plane of the phantoms, and the dominant orientation is consistent with the overall fibre arrangement in regions where fibres are arranged coherently. In the crossing phantom, where the phantom pores cross at 90°, the dominant orientation and primary dispersion orientation are assigned unpredictably between one of two directions within the printing plane (Figure 9). The secondary dispersion orientation is perpendicular to the printing plane in all voxels of all the phantoms.

## 4 Discussion

The results showed how diffusion MRI representation and model parameters respond to fibre curvature and fibre crossings in five 3D printed phantoms. A number of patterns emerge across the representations and models we investigated. Fibre curvature and crossing angle usually both affect metrics, with decreasing radius of curvature and increasing fibre crossing angle changing the metric in the same direction (but often by a different amount). We can refer to both changes as an increase in fibre orientation dispersion, as both broaden the number of directions in which water molecules can diffuse unrestricted within one voxel. Both forms of orientation dispersion also contradict the idea of a central axis of diffusion assumed by some models, and this affects the performance of those models in different ways.

### 4.1 Parametric Representations

In DTI, increased orientation dispersion of either form increases RD and decreases AD, bringing the two metrics closer together and reducing the FA. This demonstrates that axial and radial diffusivity can change significantly without any change in the underlying microstructure, and underscores the care that must be taken when interpreting those metrics in terms of tissue microstructure (Wheeler-Kingshott and Cercignani, 2009).

Fibre crossings have a stronger effect on diffusivity metrics than fibre curvature does, which is consistent with the overall lower levels of fibre dispersion for bending at the arc radii used here compared to crossing fibres. This drop in FA between straight and orthogonally crossing fibres observed in physical phantoms replicates prior simulation findings (Alexander et al., 2001), and shows a linear relationship between crossing angle and FA.

MD does not appear to systematically change with fibre crossing angle, which is expected due to the rotational invariance of MD at both the micro- and macroscopic level, but contradicts analytical findings (Vos et al., 2012). This contradiction may be resolved by observing that the variation in MD predicted by Vos et al. is small compared to the unexplained variance in MD (from noise) measured in this study. Overall, there is a larger MD for phantoms with curvature (bending, fanning, kissing) compared to phantoms with straight printing only (straight and crossing), which suggests that there is generally more free water when the printing is done in a curved manner. From Fig. 1, it is observed that the curved printing is achieved using short straight segments, which is expected to result in more gaps between printed lines compared to straight collinear printing. From our microscopy findings in previous work (Mushtaha et al., 2021), these gaps become filled with free water, which is consistent with these trends in MD. This effect would be expected to be exacerbated at higher curvature (see Fig. 1), which is consistent with the MD increases observed with increasing fibre curvature in the bending phantom.

DKI’s consideration of non-Gaussian diffusion reduces the dependence of the diffusivity metrics on orientation dispersion compared to DTI; that is, kurtosis metrics account for some of the signal variation with increased orientation dispersion. MK increases with both forms of orientation dispersion, which is expected given that mean kurtosis increases with orientation dispersion (Lasič et al., 2014). MK’s increase at low arc radii and generally higher values in the straight and crossing phantoms may also be explained by larger gaps between 3D printing lines for curved printing, similar to MD.

### 4.2 Compartment Models

The 3AM phantoms in this study essentially have two major compartments, as illustrated in Figure 1b: space within the microscopic fibrous pores, and free water between lines of extruded material. Prior work estimated the volume fraction of free water in the phantoms due to these gaps between lines to be 34% for straight-line printing, and the remaining 66% of water to be within pores (Mushtaha et al., 2021). With sufficient diffusion time for molecules within the pores to be restricted by the pore boundaries, we would expect the intra-pore space to produce diffusion signal analogous to intra-neurite space.

All three compartment models assign 20-35% volume fraction to intra-neurite space, modeled as restricted diffusion, across all five phantoms. This is considerably less volume fraction than the ∼66% volume fraction of water within pores in the phantoms, suggesting that at the diffusion times used in this experiment (∼ 13 ms), much of the diffusion within pores is not restricted. For our diffusion time, the 1D root mean squared displacement of free water at room temperature is 7.6 µm, which is the same order of magnitude as the expected median pore size of 3.5 µm and smaller than many of the larger pores. The handling of this unrestricted diffusion differentiates the ball and stick, NODDI, and Bingham-NODDI models.

Of the three compartment models, only the ball and stick model’s estimation of intra-neurite volume fraction was changed by the presence of crossing fibres. The ball and stick model has no dedicated mechanism to describe dispersing fibres, and so can only represent the decreased anisotropy caused by crossing fibres by assigning more volume fraction to the ball compartment. NODDI and Bingham-NODDI’s ODI metrics respond to crossing fibres, allowing the intra-neurite volume fractions to stay consistent.

Fibre curvature appears to have a significant effect on how the three compartment models assign volume fraction to the intra-neurite compartment. The intra-neurite volume fraction is consistently higher in the straight and crossing phantoms than in the fanning and kissing phantoms at comparable low and high crossing angles, which is consistent with the phantoms with curvature having more free water between printing lines and less restricted water in pores (as described above for MD and MK).

NODDI and Bingham-NODDI estimate the CSF volume fraction to be anywhere from 0% to about 60%. This variability in the CSF volume fraction while the intra-neurite volume fraction remains relatively consistent indicates that some signal from free water is being assigned to the extra-neurite compartment. The difficulty NODDI has distinguishing between the CSF and extra-neurite compartments in these phantoms may be explained by observing that due to NODDI’s tortuosity model, the extra-neurite compartment is modeled by a distribution of cylindrically symmetric tensors with radial diffusivity 65-75% of free water’s diffusivity. This Watson distribution of lightly anisotropic tensors generates a diffusion pattern that is similar to isotropic diffusion, which may lead to an unstable fit between the extra-neurite and CSF compartments. A phantom with a more structurally complex microstructure including a third compartment, or a scan protocol with a longer diffusion time resulting in more restricted diffusion would perhaps elicit a more consistent response from NODDI.

NODDI’s volume fraction metrics have limited sensitivity to the orientation dispersion presented by these phantoms, which suggests that changes in NODDI’s volume fraction metrics indicate changes to the microstructural fibre composition, as intended. NODDI’s ODI metric changes reliably with fibre crossing angle, suggesting that it may be a robust indicator of orientation dispersion, even when there are volume fraction instabilities. However, the planar patterns of orientation dispersion in the 3AM phantoms challenge the Watson distribution: to distribute the intra-neurite compartment’s orientation along a plane, it needs to distribute it equally perpendicular to that plane. There is an inherent trade-off between the accuracy of the predicted diffusion pattern in-plane and out-of-plane. This trade-off forced by NODDI’s use of the Watson distribution is addressed by Bingham-NODDI.

While Bingham-NODDI also has a somewhat unstable fit of volume fractions, it is more stable than NODDI (qualitatively and as evidenced by higher R^2^) and orientation dispersion affects the volume fraction Bingham-NODDI assigns to the CSF and extra-neurite compartments. Regions with a lower crossing are more often assigned no volume fraction of CSF, especially in the fanning phantom. The relatively low arc radii in the bending phantom data also demonstrate Bingham-NODDI’s tendency to assign no volume fraction to the CSF compartment, which is consistent with the observation that regions with a low crossing angle in the fanning phantom also tend to have a lower arc radius than other regions (Fig. 1). These observations are also consistent with the phantoms containing more free water for higher curvatures.

The high sensitivity of ODI_P_ to fibre crossing angle demonstrates that the Bingham distribution is able to represent the planar patterns of diffusion observed in the 3AM phantoms without the tradeoffs forced by the Watson distribution. This is further underscored by the total lack of dispersion perpendicular to the printing plane indicated by Bingham-NODDI. However, the phantoms used in this study are limited to planar patterns of dispersion, so we could not compare the performance of NODDI and Bingham-NODDI in the presence of 3D patterns of dispersion. ODI_P_ spans its entire defined range in the fanning and kissing phantoms, granting this metric high sensitivity to crossing fibres.

This work illustrates a physical dMRI phantom that produces a biophysically plausible ground truth for diffusion signal in the presence of complex fibre configurations like fanning and kissing fibres. While analyses similar to those presented here were previously available using simulated data (Alexander et al., 2001), the level of orientational complexity easily achievable with physical phantoms was more limited. The 3D printed phantoms therefore fill a previously empty niche in the spectrum of phantoms available for the validation of dMRI, offering a low cost and accessible 3D printing approach with greater control over the ground truth fibre geometry compared to ex-vivo tissue and some existing physical phantoms, along with the ability to generate scan data over a range of complex orientations without the computational requirements of a numerical phantom.

### 4.3 Limitations

The nature of the 3D printing technique used to produce the phantoms left some artifacts in the phantom. Some air bubbles were trapped in the phantoms, leading to round, dark areas in the b0 and diffusion-weighted images. These artifacts are readily masked out, but reduce the amount of usable data from each phantom. Arcs are printed in short, straight segments, and those segments are long enough to be individually visible in the shorter arcs. In these cases, the printed pattern is a poor approximation of an arc. This also resulted in free water partial volume that varied with curvature, which needs to be considered when interpreting results; however, this effect was small compared to the impact of crossing fibre angle on dMRI parameters. The layer-by-layer 3D printing procedure also means that the complex geometries are restricted to a single plane. In other words, the diffusion parallel to the axis of the cylindrical phantoms is constant, regardless of the in-plane print pattern, which limits the geometric complexity achievable with the 3D printed phantoms.

Underextrusion of filament during the 3D print process led to unintended gaps between lines of material, and a relatively high volume fraction of free water throughout the phantoms that varied with printing curvature. While this structure was practically useful because it allowed water to reach and dissolve pockets of PVA in the elastomeric matrix more easily, it also resulted in a less anatomically realistic microstructure than there would be with less free water.

Producing very small radii of curvature with a 3D printer requires a high print resolution. At the resolution used by this iteration of the CURA extension, the smallest circle of material in the phantom consisted of only three straight segments (as shown in the centre of the arcs in Figure 1a), likely allowing a higher volume fraction of water in those regions. The potentially higher volume fraction of free water in regions with a small arc radius challenges interpretation of results in these regions.

The diffusion time used in these experiments was low compared to typical values used for *in vivo* diffusion MRI in the human brain. While this may be somewhat justified by the lower mean-squared displacements of water expected at room temperature compared to body temperature, exploring the diffusion time dependence of these models is beyond the scope of this work that aims to introduce this new class of crossing fibre phantoms and demonstrate their potential utility for several common models. Furthermore, the relatively low diffusion time prevents some of the larger pores in the phantom from restricting diffusion, contributing to NODDI and Bingham-NODDI’s unstable fit between the extra-neurite and CSF compartments.

### 4.4 Representation and Model Performance

Using these phantoms, we can identify a set of representation and model parameters with the potential to have particular utility due to invariance or sensitivity to orientation dispersion. Specifically, MD in DTI and intra-neurite volume fraction in both NODDI and Bingham-NODDI were largely invariant to fibre crossing angle. This invariance means that they are potentially robust indicators of microstructural change in the presence of multiple fibres. Meanwhile, ODI in NODDI and ODI_P_ in Bingham-NODDI both have strong relationships with fibre crossing angle, which suggests that they may be good indicators of orientation dispersion. These results also show potential overfitting between NODDI and Bingham-NODDI’s extra-neurite and CSF compartments. This overfitting may be caused by the fixed CSF diffusivity matching the axial diffusivity in the other compartments.

### 4.5 Conclusions

This work illustrates a physical dMRI phantom that produces a biophysically plausible ground truth for diffusion signal in the presence of complex fibre configurations like fanning and kissing fibres. While analyses similar to those presented here were previously available using simulated data (Alexander et al., 2001), the level of orientational complexity easily achievable with physical phantoms was more limited. The 3D printed phantoms therefore fill a previously empty niche in the spectrum of phantoms available for the validation of dMRI, offering a low cost and accessible 3D printing approach with greater control over the ground truth fibre geometry compared to ex-vivo tissue and some existing physical phantoms, along with the ability to generate scan data over a range of complex orientations without the computational requirements of a numerical phantom.

## Author Contributions

TK designed the phantoms, wrote the software to produce the phantoms, helped with the phantom production process, and wrote the analysis code. FM contributed to the study design and phantom production process. AK and CB contributed to the study design and supervised the execution of the study and analysis.

## Acknowledgements

This work previously appeared online in a Masters thesis (Kuehn, 2020).

## Data Availability

The Ultimaker Cura extension used to produce the phantoms is available on GitHub [https://github.com/tkkuehn/CuraEngine]. The dataset generated for this study can be found on OSF [https://osf.io/4cdw3/]. The code used to analyze that dataset can be found on GitHub [https://github.com/tkkuehn/dmri-phantom-applications].

